# The relative values of the turnover number and the dissociation rate constant determine the definition of the Michaelis-constant

**DOI:** 10.1101/052514

**Authors:** Gassan Nazzal

**Keywords:** K_m_ value, turnover number, Rapid equilibrium approximation, diffusion limited reaction, enzymatic efficiency, Transition state affinity

## Abstract

In this work, we attempt to determine the assumptions of each case of the QSSA. We came to the conclusion that for an enzyme with average kinetics parameters the REA is a good approximation to derive the rate equation and the Km value tends to equal the dissociation constant K_d_. The active site classifies the population of the substrate into two energy states, the ground state, and the transition state. The ratio K_m_/K_d_ is equal to the partition function of the assumed two-state-system. For the average enzyme, the partition function of the transition tends to equal 1 thus the majority of the substrate molecules are in the ground state and the assumption kcat << k_−1_ is valid hence Km ≈ Kd. In contrast, when the enzyme is diffusion controlled, the Km value is equal to the productive dissociation rate k_cat_/k_1_. We have also redefined the Km value as the equivalence point of the reaction rates, namely, the effective diffusion rate and the maximal catalytic rate, which reflects more clearly the transition from the bimolecular reaction to the unimolecular reaction in the saturation curve.

## 1. Introduction

Michaelis and Menten applied the rapid equilibrium approximation (REA) on the hydrolysis of sucrose by the enzyme Invertase, in the first step the enzyme binds the substrate reversibly, in the second step, the irreversible reaction takes place ^[1]^

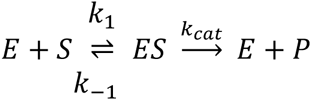
k_1_: The association rate constant of ES – complex (M^−1^s^−1^). k_−1_: The unproductive dissociation rate constant of ES – complex (s^−1^). k_cat_: The turnover number (s^−1^).

The REA states that the dissociation rate constant k_−1_ is fast in comparison with the turnover number k_cat_. Thus, we can assume that the substrate concentration at the half maximal velocity is equal to the dissociation constant K_d_, which indicates the affinity between E and S. This assumption is a special case of the quasi-steady-state approximation (QSSA) proposed by G. Briggs and J. Haldane in 1925 ^[2]^. The QSSA defines the K_m_ value as:

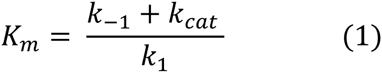

The REA is valid when k_−1_ >> k_cat_:

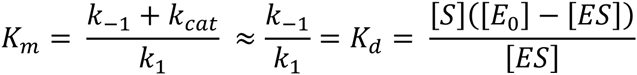

The second special case of the QSSA is the Van Slyke-Cullen mechanism^[3]^, where both steps are irreversible and k_−1_ << k_cat_:

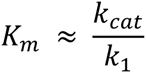

The efficiency of the enzyme provides information about the ratio between k_−1_ and k_cat_. If we assume that the association step is diffusion controlled thus we can assume that the association rate constant k_1_ is almost equal to the diffusion rate of the molecules and The enzyme efficiency can be determined by comparing the k_cat_/K_m_ value with the diffusion limit of a bimolecular reaction.:

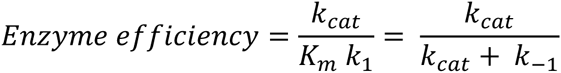

In this work, we will determine the assumptions of each case of the QSSA. To do so we have to study the reaction rate constants, that defines the K_m-_ value to estimate an equation that predicts the enzyme efficiency.

## 2. Kinetics interpretation of the saturation behavior

Before the reaction takes place the molecules have to diffuse and collide with each other and only the collisions in the right point on the surface of the enzyme and with the sufficient amount of energy will lead to the formation of the product. The sequence of one enzymatic cycle can be divided into two steps, namely the effective diffusion time to form an activated ES-complex and the duration of the maximal catalytic rate:

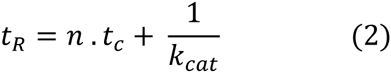
t_R_: the duration of one enzymatic cycle. n: denotes the total number of S molecules that are needed to hit the active site every cycle, until an energetically active molecule binds to form a product. We will define this parameter as the collision number. t_C_: diffusion-collision time, time until E and S encounter each other.

Assuming that all molecules are mixed and distributed equally in the solution, we can calculate the needed time to form an ES complex as a function of the substrate concentration, using the mean squared displacement equation. Firstly we define the diffusion-collision time as:

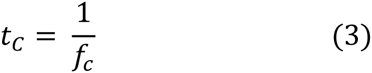

The collision frequency can be expressed as follows:

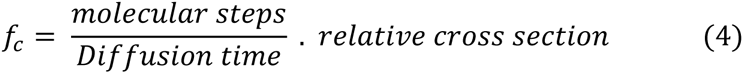

We define one molecular step as the distance at which the molecule translocates its own diameter (2r). To diffuse a distance x, the molecule had to diffuse x/2r molecular steps, and for two molecules the number of steps will be:

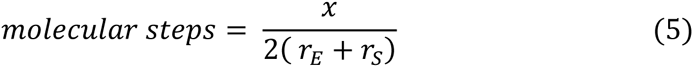

The relative reaction cross section is the ratio between the targets area of the collision event and the total available area. In this case:

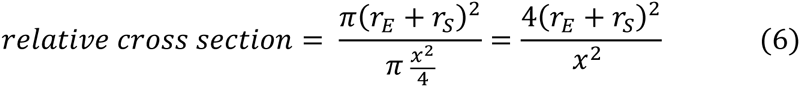

To estimate the diffusion distance x, we assume that the substrate molecules in the solution are equally distributed in the total volume V. Thus, each substrate molecule occupies the volume V/N_S_ (divided by 1000 to convert from liter to m^3^), N_A_ denotes the Avogadro’s number:

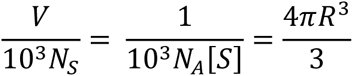

We define the diffusion distance as the diameter of the spherical volume per substrate molecule:

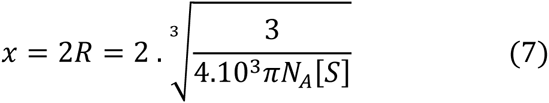

To calculate the diffusion time for two-body problem, we take the sum of the diffusion velocities of both molecules E and S, using the equation of the mean squared displacement for a three-dimensional case (t = x^2^ / 6D):

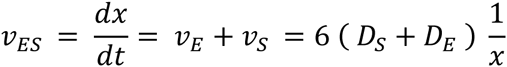

We assume that the substrate molecule diffuse the total volume per molecule until the collision occurs thus we can estimate the integral with the following boundary conditions to calculate the diffusion time:

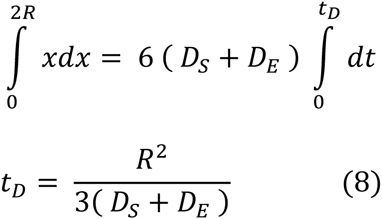

Substituting eq (5), (6), (7) and (8) in eq (4) we obtain:

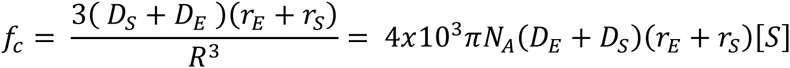

Substituting the Stokes-Einstein– relation in the last equation we obtain equation (9):

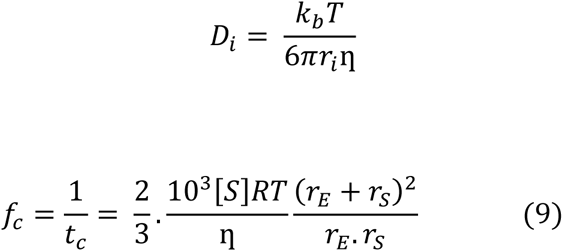
f_c_: the collision frequency at a given substrate concentration (s-^1^). η: denotes the dynamic viscosity (N.s.m-^2^). r_E_ and r_s_ the radius of each molecule (m). [S] the substrate concentration (mol/l = M).

One of the limitations of the equation (9) is that the substrate concentration will decrease with the time as a result of the enzymatic activity. Consequently, the collision frequency is time-dependent. Therefore, we must adjust that [S]_0_ >>[E]_0_ to assume the free ligand approximation [S]t ≈ [S]_0_.

The reaction rate can be calculated by multiplying the catalytic frequency f_cat_ (s^−1^) of one active site and the total enzyme concentration in molar units [E_0_]:

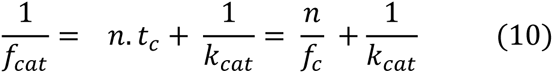

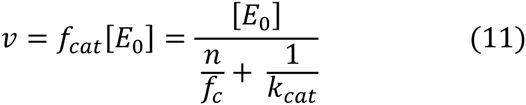

Rearranging the Michaelis-Menten equation in a similar form as equation (11), we obtain:

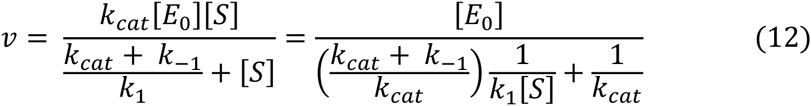

Equating equation (11) and (12):

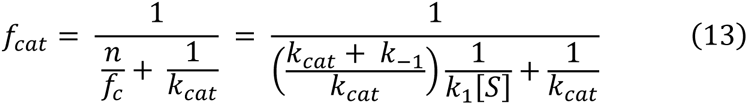

Equation (14) represents a redefinition of the Km as the equivalence point between the effective diffusion rate and the maximal reaction rate, rather than the classical concept of the enzyme saturation that describes a general protein-ligand interaction.

At [S] = K_m_, the effective diffusion rate, and the maximal reaction rate are equivalent and we obtain:

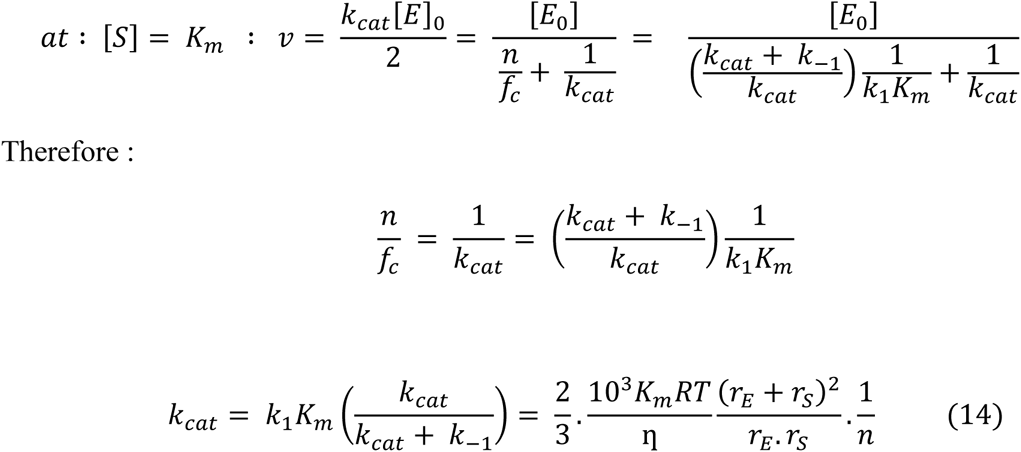

The diffusion controlled bimolecular rate constant of the ES-complex formation can be theoretically calculated as:

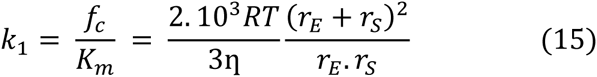

The total maximal collision frequency when [S] = K_m_ is equal to:

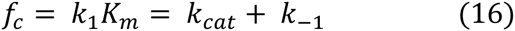

For an activation controlled enzyme the saturation behavior can be described in kinetics terms:

[*S*] < *K*_m_ *The reaction is under diffusion control thus* [*E*] > [*ES*]

[*S*] < *K*_m_ *The reaction is under diffusion control thus* [*E*] > [*ES*]

## 3. The energy profile of the enzyme action

The K_m_ value is the sum of the dissociation constant K_d_ and the effective substrate concentration Mathematical expression of this statement can be written through substituting equations (14) and (15) in equation (18) and then in equation (1) we obtain the equation (19):

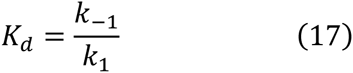

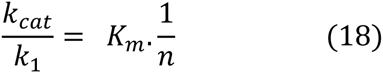

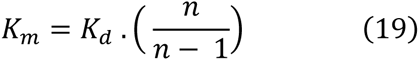

### 3.1 The collision number determines the probability of the reaction

Rearranging the K_m_ expression into two fractions allow us to treat the system as a canonical ensemble of the ES-complex. The two fractions denote the probabilities of two dissociation events, namely, the event that the complex will stay in association-dissociation-equilibrium with the free substrate and the free enzyme and the event that the complex will reach the transition state and react:

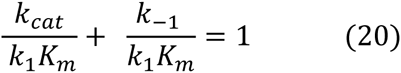

Enzymes accelerate the reaction rate through destabilizing the substrate structure by a certain reaction mechanism, increasing in this manner the probability that the substrate will reach the transition state. The Boltzmann distribution allows us to relate the fraction of molecules in a certain state to the probability to find this state:

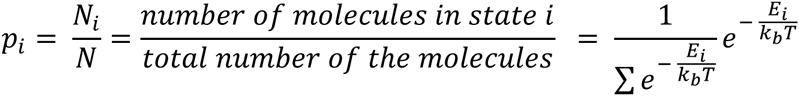

P_i_: the probability of the state i in the system.

E_i_: The energy level of the state i. k_b_: The Boltzmann constant, T: The absolute temperature.

In one catalytic cycle, one substrate molecule will be converted to a product. In one second the productivity is equal to the turnover number which indicates the successful dissociation of the ES-complex after reaching the transition state. The probability p_‡_ to find an ES-complex in the transition state with the energy level E‡ is:

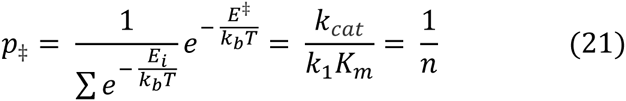

On the other hand (n - 1) ES molecules are in equilibrium with the free form of the enzyme and the substrate. Assuming that the association event is diffusion controlled, the dissociation event is under thermodynamic control and had to overcome the dissociation energy. The probability p_0_ to find an ES-complex in the unproductive dissociated form is:

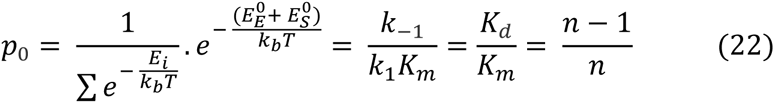

To simplify the calculations we use the activation energy ΔE^‡^, and the dissociation energy of the ES-complex ΔE_diss_, E^0^_ES_ denotes the ground state of the ES-complex:

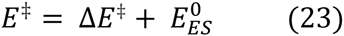

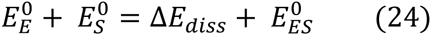

Substituting eq(23) in eq(21) and eq(24) in eq(22) and then estimating the ratio p_‡_/p_0_ to determine the collision number, we obtain the following expression of the collision number:

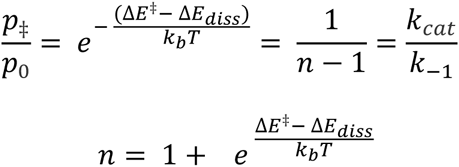

Thus, the enzyme efficiency is the probability that the ES-complex will reach the transition state. The probability of the catalytic event and the collision number are determined by the difference between the dissociation energy barrier and the activation energy barrier:

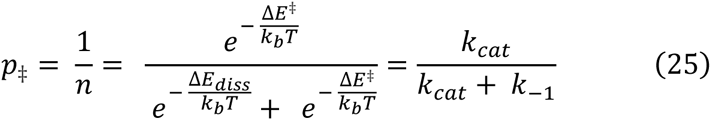

When ΔE^‡^> > ΔE_diss_, then we obtain the expression of the Arrhenius equation of the activation energy E_a_:

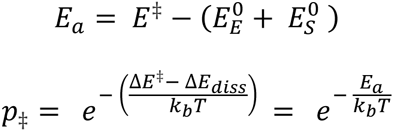

The ratio between K_d_ / K_m_ is the probability that the ES-complex is in association–dissociation equilibrium:

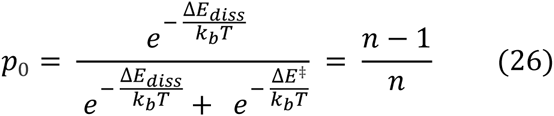

The ratio K_m_ / K_d_ can be evaluated as a vibrational partition function relative to the ground states of the free enzyme and the free substrate:

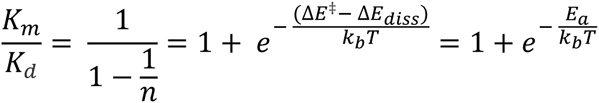

The evaluated partition function represents the ability of the catalytic reaction mechanism to destabilize the ground state of the substrate, populating in this way the transition state. When the partition function tends to equal 1, this means that the majority of the ES-molecules exist in the ground state and the active site exhibits a low catalytic activity, therefore the system exists in association - dissociation equilibrium K_m_ = K_d_.

### 3.2 The affinity toward the transition state determines the probability of the reaction

Using the Eyring-equation we can express equation (1) in terms of free energy. For zero order reaction the relation between the rate constant and the free energy of the activation ΔG^‡^ is (assuming that the transmission coefficient equals 1):

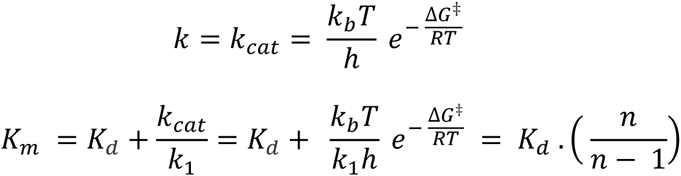

Substituting the relation between the dissociation constant and the free energy of the binding ΔG_b_ we obtain:

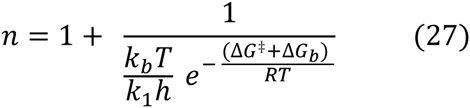

We can now express the probability of the catalytic event in terms of free energies using equation (27)

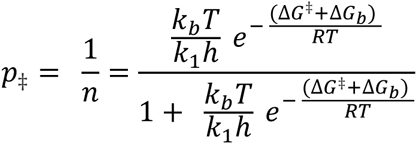

The Eyring theory assumes that the reactant and the transition state are in rapid equilibrium state, thus, we can express the equilibrium constant of the transition as:

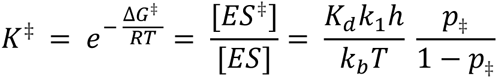

The ratio K^‡^/ K_d_ represents the affinity of the enzyme toward the transition state:

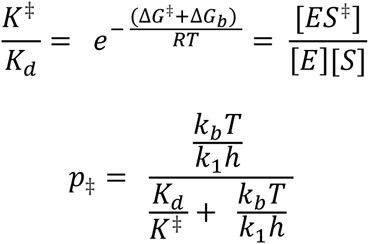

The ratio K^‡^ / K_d_ represents the ability of the active site firstly to recognize the right substrate and secondly to destabilize it. The sum of the free energy differences of both steps indicates that the two enzymatic steps are coupled, the binding step is exergonic while the destabilizing step is endergonic. thus, we can assume that the substrate binding induces a conformational change that is complementary to the transition state. The probability to induce a conformational change that opens the reaction path is maximized when the sum of the free energies differences of the ground state and the transition state formation is minimized. This is a conformational proofreading mechanism ^[4]^, the substrate binding induces a conformational change that causes a deformation in the substrate structure, hence, the lower the affinity toward the substrate the lower the probability to induce the transition state.

## 4. Defining the assumptions of each case of the QSSA

1-Activation controlled enzyme: The energy difference between the activation energy and the dissociation energy is high, thus the catalytic event is unlikely to proceed and the active site exhibits a low stabilizing ability of the transition state. The majority of the association events dissociate unproductively and the system exists in a rapid equilibrium state:

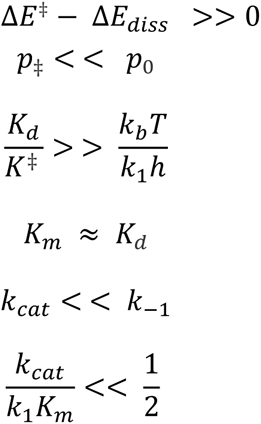

### The validity of the REA

The average enzyme exhibits a k_cat_ of 10 s^−1^ and a K_m_ value of 100 μM ^[5]^. Taking the theoretical diffusion limit as 10^9^ M^−1^s^−1^ to calculate the ratio (k_cat_/ K_m_) / k_1_ will yield an enzyme efficiency of 0.01%, which is equal to the relative deviation between K_m_ and K_d_, thus, the REA is valid on enzymes with average kinetics parameters.

**Figure 1.**
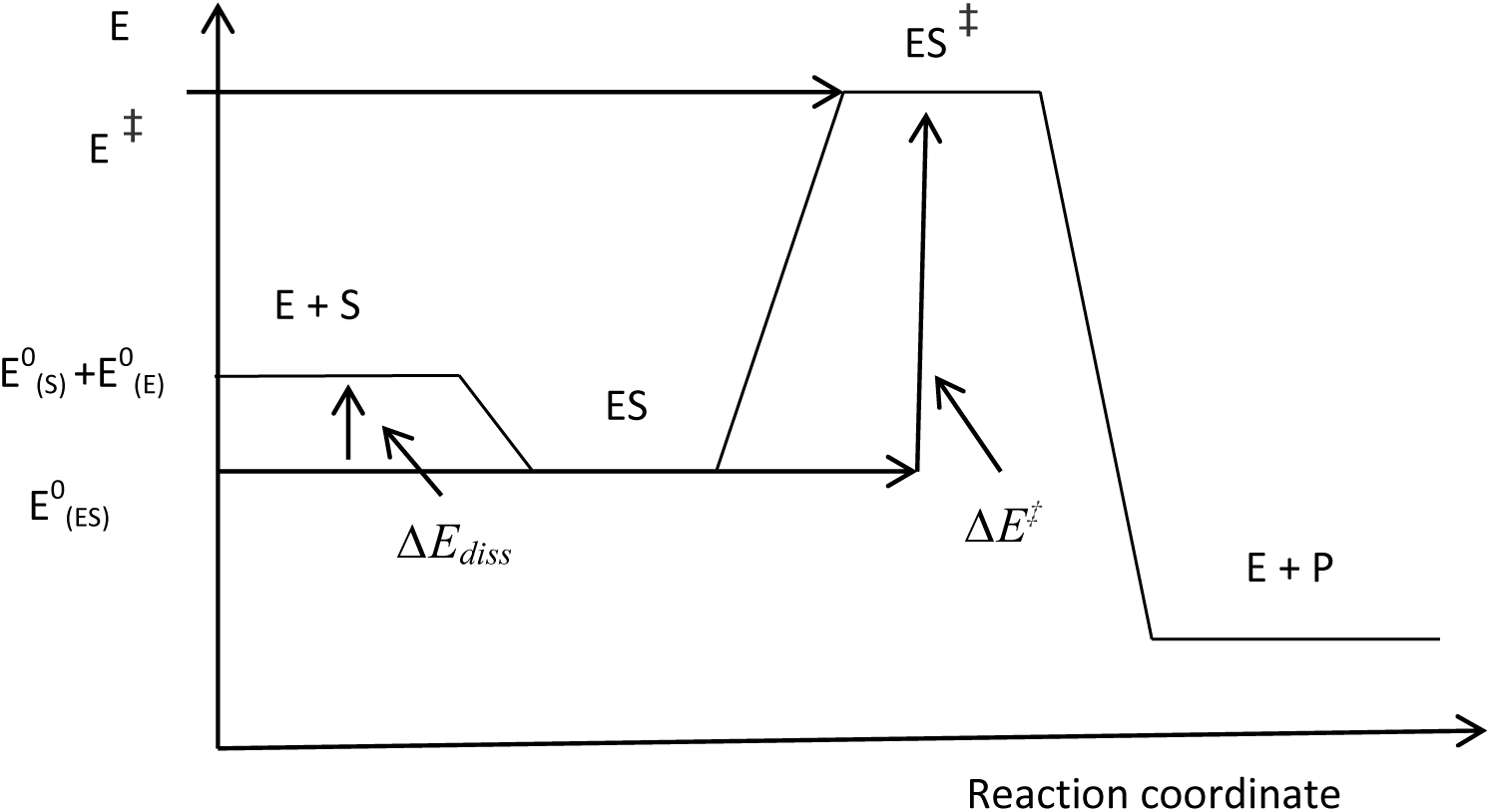
***Case 1*** the difference between the activation energy and the dissociation energy is high thus the majority of the molecules are in rapid association-dissociation equilibrium.

2- The energy difference between the activation energy and the dissociation energy equals zero, therefore, the ES-complex dissociates equally in both directions and the enzymatic efficiency is equal to 50% of the effective collisions. The ES-complex will dissociate in both directions equally and the reaction is diffusion controlled:

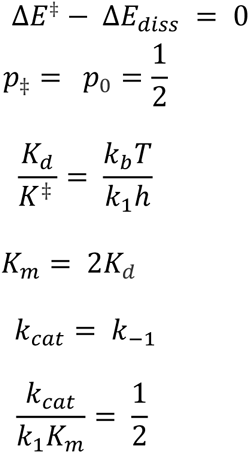

**Figure 2.**
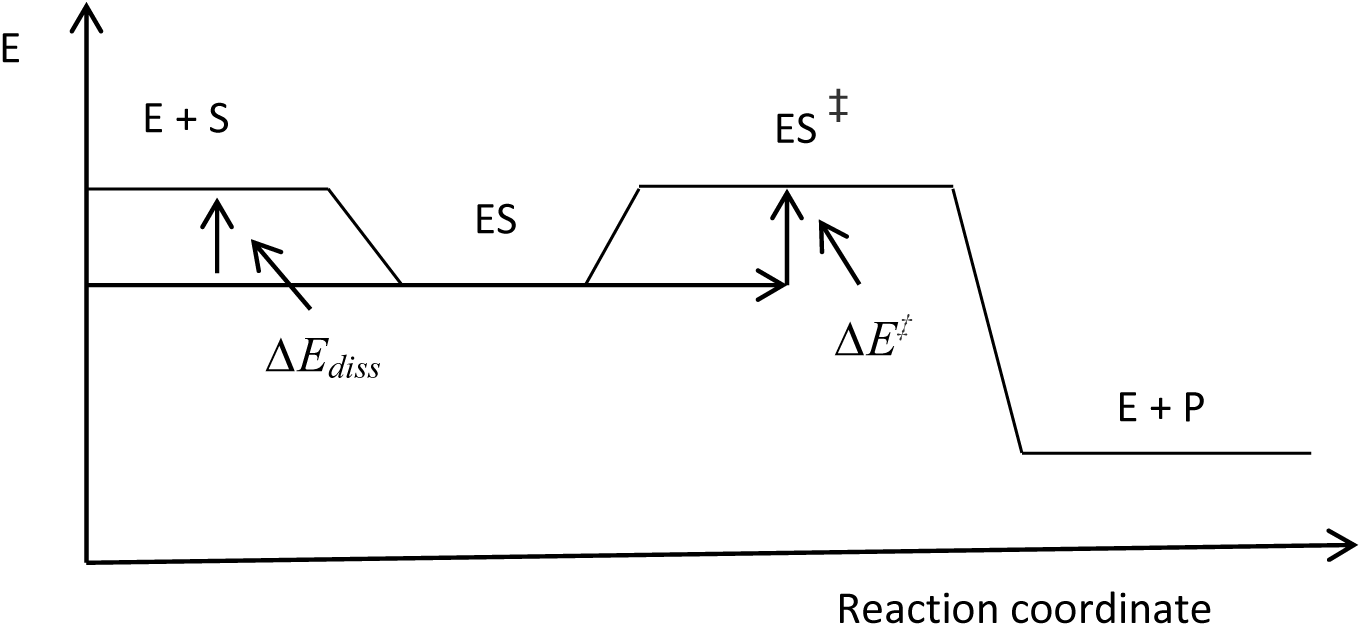
***case 2*** the activation energy and the dissociation energy are equal thus, the probabilities of the dissociation are equally probable in both directions.

3- Diffusion-controlled enzyme: The activation energy barrier is lower than the dissociation energy thus the dissociation is directed forward irreversibly in a way that nearly every collision leads to a reaction:

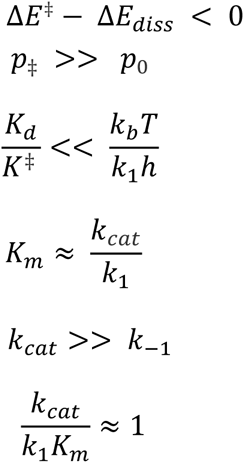

The difference between the activation energy and the dissociation energy becomes negative if the reaction proceeds stepwise each step had a relatively low activation energy barrier to overcome and the released energy from the first reaction step makes the total difference ΔΔE more negative.

## 5. The productive dissociation constant k_cat_/k_1_

In the literature, the value of k_cat_/K_m_ is used as a measure of the enzyme efficiency because it can be easily compared with the diffusion controlled association rate constant k_1_ of a bimolecular reaction. In contrast to k_cat_/K_m_ the ratio k_cat_/k_1_ represents the substrate concentration that could be converted to products at half-maximal reaction rate:

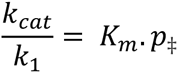

To understand the reason why some enzymes work kinetically perfect and others do not, we need to consider the biological context of the enzyme kinetics.

An activation controlled enzyme may be kinetically inefficient but in vivo, this inefficiency may be very important for the cellular homeostasis and the biochemical control. Assuming that the ratio k_cat_/k_1_ is the wanted physiological concentration of the product in the cell, increasing the efficiency will lead to an accumulation of the product that may distort the homeostasis, which leads to a pathological state. Another reason for a low efficiency is, that the substrate consumption may be shared by several pathways, therefore, increasing the efficiency of one enzyme in one pathway will lead to a higher consumption of the substrate in the kinetically more efficient pathway which may cause a deficiency in the product formation in the less efficient pathway. Hypothetically if every enzyme in the cell would be diffusion controlled the organism will need additional regulating mechanisms to maintain the concentrations of the products and the substrates in the physiological range, because the enzyme accelerates the achievement of the equilibrium state, which is not the physiological state in most cases. Thus, the pre-defined enzyme efficiency will yield a defined product formation without the need to add more regulatory points, which will result in a more complex biochemical map. Hence, the activation energy from the biological perspective is a required protective barrier to maintain the stability of the cellular structure and function.

A high efficiency would be beneficial in the elimination of toxic compounds to protect the cell. A diffusion-controlled enzyme will eliminate a specific cytotoxic compound in every encounter, typical examples are: catalase, superoxide dismutase, β-lactamase ^[7]^. Another biological function that needs a total elimination of the substrate would be the switch of a biological function between on or off mode, a typical example is the acetylcholine-esterase. An accelerated total elimination of the neurotransmitter from the synaptic cleft is essential in decreasing the regeneration time, making the system in this way ready to sense a new signal.

**Table 1:** Theoretical calculations: we have maximized theoretically the enzyme efficiency p_*‡*_ at constant K_m_ value, K_m_ = 100 μM. The diffusion limit is calculated by assuming that the substrate and the enzyme are in the same size r_E_ = r_S_, temperature at 25 °C, and the dynamic viscosity is equal to 1 mPa.s.

| $p_{\ddagger}$ | 0.99 | 0.8 | 0.5 | 0.2 | $10^{-1}$ | $10^{-2}$ | $10^{-4}$ | $10^{-6}$ |
| --- | --- | --- | --- | --- | --- | --- | --- | --- |
| n | 1.01 | 1.25 | 2 | 5 | 10 | 100 | $10^4$ | $10^6$ |
| $\Delta\Delta E$<br>kJ/mol | -11.382 | -3.433 | 0 | 3.433 | 5.442 | 11.38 | 22.813 | 34.22 |
| $K_d$<br>$\mu M$ | 1 | 20 | 50 | 80 | 90 | 99 | 99.99 | 99.999 |
| $K^{\ddagger}$ | $1.04 \times 10^{-7}$ | $8.48 \times 10^{-8}$ | $5.3 \times 10^{-8}$ | $2.12 \times 10^{-8}$ | $1.06 \times 10^{-8}$ | $1.06 \times 10^{-9}$ | $1.06 \times 10^{-11}$ | $1.06 \times 10^{-13}$ |
| $K^{\ddagger}/K_d$<br>$M^{-1}$ | 0.104 | $4.24 \times 10^{-3}$ | $1.06 \times 10^{-3}$ | $2.65 \times 10^{-4}$ | $1.17 \times 10^{-4}$ | $1.07 \times 10^{-5}$ | $1.06 \times 10^{-7}$ | $1.05 \times 10^{-9}$ |
| $\Delta G_b$<br>kJ/mol | -34.221 | -26.800 | -24.530 | -23.366 | -23.074 | -22.838 | -22.814 | -22.814 |
| $\Delta G^{\ddagger}$<br>kJ/mol | 39.827 | 40.332 | 41.497 | 43.766 | 45.483 | 51.187 | 62.594 | 74.024 |
| $k_{cat}/k_1$<br>$\mu M$ | 98 | 80 | 50 | 20 | 10 | 1 | 0.01 | 0.0001 |
| $k_{-1}$<br>$s^{-1}$ | $6.6 \times 10^3$ | $1.32 \times 10^5$ | $3.3 \times 10^5$ | $5.28 \times 10^5$ | $5.49 \times 10^5$ | $6.53 \times 10^5$ | $6.59 \times 10^5$ | $6.59 \times 10^5$ |
| $k_{cat}$<br>$s^{-1}$ | $6.53 \times 10^5$ | $5.28 \times 10^5$ | $3.3 \times 10^5$ | $1.32 \times 10^5$ | $6.6 \times 10^4$ | $6.6 \times 10^3$ | 66 | 0.66 |
| $k_{cat}/K_m$<br>$M^{-1}s^{-1}$ | $6.53 \times 10^9$ | $5.28 \times 10^9$ | $3.3 \times 10^9$ | $1.32 \times 10^9$ | $6.6 \times 10^8$ | $6.6 \times 10^7$ | $6.6 \times 10^5$ | $6.6 \times 10^3$ |

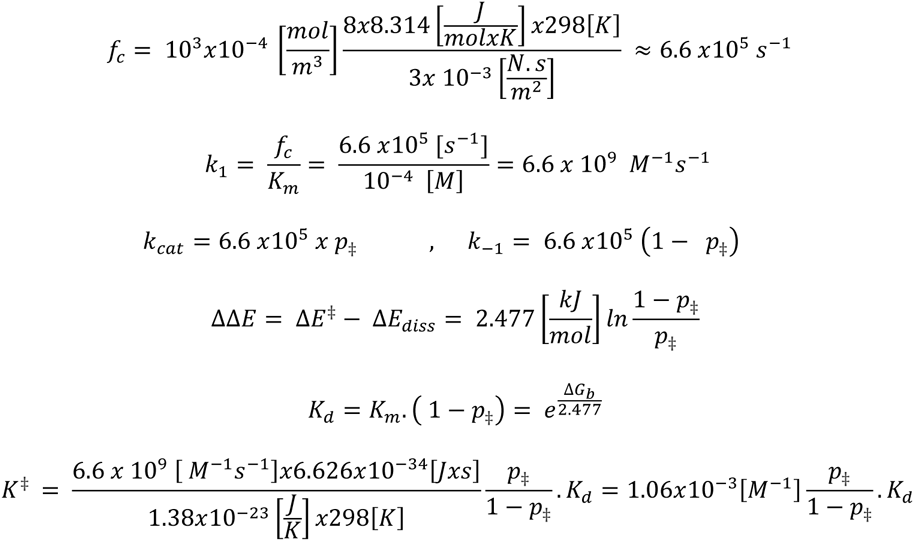

## References

[1] Michaelis L, Menten ML. “Die Kinetik der Invertinwirkung”. Biochemische Zeitschrift. 1913, 49: 333–369.

[2] Briggs, G.E.; Haldane, J.B.S. (1925). “A note on the kinetics of enzyme action”. Biochem J 19 (2): 338–339.

[3] K Brocklehurst and A Cornish-Bowden, “The pre-eminence of k_cat_ in the manifestation of optimal enzymic activity delineated by using the Briggs-Haldane two-step irreversible kinetic model”, Biochem J. 1976 Oct 1; 159(1): 165–166.

[4] Savir Y, Tlusty T (2007) “Conformational Proofreading: The Impact of Conformational Changes on the Specificity of Molecular Recognition”. PLoS ONE 2(5): e468. doi:10.1371/journal.pone.0000468

[5] Bar-Even, A; Noor, E; Savir, Y; Liebermeister, W; Davidi, D; Tawfik, DS; Milo, R. “The moderately efficient enzyme: evolutionary and physicochemical trends shaping enzyme parameters”. Biochemistry, May 31, 2011 50 (21): 4402–10.

[6] Berg JM, Tymoczko JL, Stryer L, Biochemistry. 5th edition: p. 329

